# Sex-specific transgenerational effects of diet on offspring life history and physiology

**DOI:** 10.1101/2022.05.23.492998

**Authors:** Tara-Lyn Camilleri, Matthew D.W. Piper, Rebecca L. Robker, Damian K. Dowling

## Abstract

Dietary variation in males and females can shape the expression of offspring life histories and physiology. However, the relative contributions of maternal and paternal dietary variation to phenotypic expression of latter generations is currently unknown. We provided male and female *Drosophila melanogaster* diets differing in sucrose concentration prior to reproduction, and similarly subjected grandoffspring to the same treatments. We then investigated the phenotypic consequences of this dietary variation among grandsons and granddaughters. We demonstrate transgenerational effects of dietary sucrose, mediated through the grandmaternal lineage, which mimic the direct effects of sucrose on lifespan, with opposing patterns across sexes; low sucrose increased female, but decreased male, lifespan. Dietary mismatching of grandoffspring-grandparent diets increased lifespan and reproductive success, and moderated triglyceride levels, of grandoffspring, providing insights into the physiological underpinnings of the complex transgenerational effects on life histories.

## Main

From nematodes to primates, parental environments may shape the phenotypes of their offspring through non-genetic mechanisms that are either condition dependent or epigenetic in origin^1–3 4^. Consequently, when individuals are subjected to environmental heterogeneity prior to reproduction, their exposure to these environments can shape components of fitness in offspring and subsequent generations (transgenerational effects)^5–9^. Recent experiments have shown that variation in environmental factors, such as predation risk and levels of sexual conflict, among parents may catalyse transgenerational effects that differ in magnitude or direction across sexes, and which may also be lineage (genotype) specific^10^. Notwithstanding, currently it remains unclear whether such transgenerational effects are consistently instigated across diverse environmental stresses, whether they generally act to enhance or depress offspring performance, and whether they are transferred primarily through maternal or paternal lineages or hinge on interactions between both.

Nutrition is a pervasive and critical source of environmental variation that shapes phenotype. Variation in macronutrient balance or caloric content has been shown to confer direct effects on lifespan, fecundity, and underlying physiology^11–14^. Studies from diverse species have demonstrated that females and males require different diets to maximise their fitness^15–19^. Female fitness is maximised on a higher relative protein concentration because high protein facilitates egg production, while higher relative carbohydrate content for males provides fuel for attracting and locating a mate^20–24^. Recent studies have also shown dietary-induced intergenerational effects across a variety of species; for example, changes to sugar content of the parental diets in fruit flies^25,26^ or dietary fat content in mice^27^ induces phenotypic changes in parents that are transmitted to their offspring. Intriguingly, when the sucrose content of both male and female parents are altered, then parental contributions to offspring phenotypes may involve complex dam-by-sire interactions that are non-cumulative and dependent upon the sucrose content of the offspring diet ^28^. It is less clear, however, whether these dietary-mediated parental effects are epigenetic in origin, and thus inherited across multiple generations ^10,25,29–32^, and if so, whether phenotypic consequences for males and females are divergent.

Here, we experimentally tested the capacity for dietary sucrose variation among male and female fruit flies (*Drosophila melanogaster*) to precipitate transgenerational effects on components of life-history and physiology in their grandoffspring. Flies were administered one of two diets that varied in the concentration of sucrose (2.5% or 20% sucrose). The diets were administered using a full factorial design: males and females were each assigned to one of the two diets prior to reproduction, and then their grandsons and granddaughters were administered the same dietary treatments. All male-female-grandoffspring dietary combinations were represented, resulting in female-male and grandparent-grandoffspring diet combinations that were either matched or mismatched (Figure 1, panels A and B). This design enabled us to test whether dietary-mediated transgenerational effects exist, to decipher the relative grandmaternal and grandpaternal contributions, and the capacity for interactions between grandparental diets and those of the grandoffspring to shape grandoffspring phenotype, and to determine whether such effects are sex specific.

**Figure 1.**
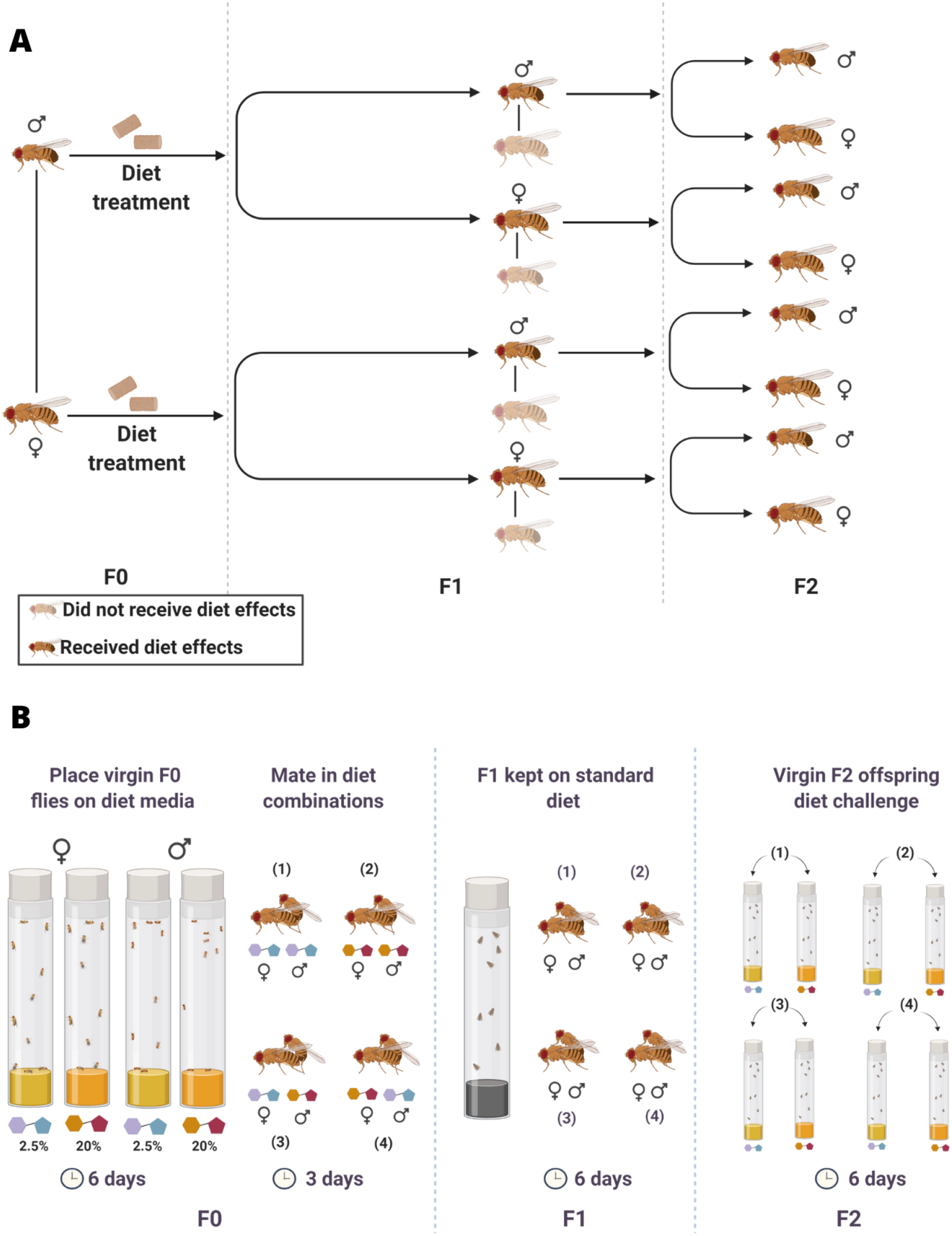
**A) Diet effects lineage.** Diet treatments were administered to both parents in the F0; and they were mated to create the F1 offspring, and received a standard diet (both males and females for each parent), and F1 offspring were mated with flies outside of the experiment that received a standard diet. This allowed us to track which F1 sex was passing on the diet effects to the F2 generation. **B) Experimental design.** The F0 generation was administered either higher (20% of overall solution) or lower (2.5%) relative sucrose in adulthood, and kept on this diet in sex-specific cohorts for 6 days as virgins before a subsequent three day cohabitation (on common garden media) that allowed mating to occur. Male and female F0 flies were combined in all possible diet combinations. The F1 generation was reared, maintained (6 days again), and cohabited (3 days) on common garden media (an intermediate sucrose content of 5%). The F2 generation was reared from egg-to-adulthood on common garden media, and then challenged as virgins with either the higher or lower sucrose such that their diet either matched or mismatched one or both of their grandparents (F0).

## Results

### Direct and indirect effects of dietary sucrose on grandoffspring lifespan are sex-specific

The diets of the grandoffspring (F2) flies conferred direct and sex-specific effects on their lifespan (*F*_1,148_ = 369.80, *p* <0.001, Table S2, Figure 2). Female F2 flies assigned to the low sucrose diet lived longer than females or males assigned to any other treatment, and 30% longer than females on the high sucrose diet. Females assigned to a high sucrose diet exhibited the shortest lifespan of any group of flies. In contrast to the large negative effect of high sucrose on female lifespan, high dietary sucrose conferred a moderate increase in male lifespan relative to males assigned to a low sucrose diet (Table S2; Figure 2).

**Figure 2.**
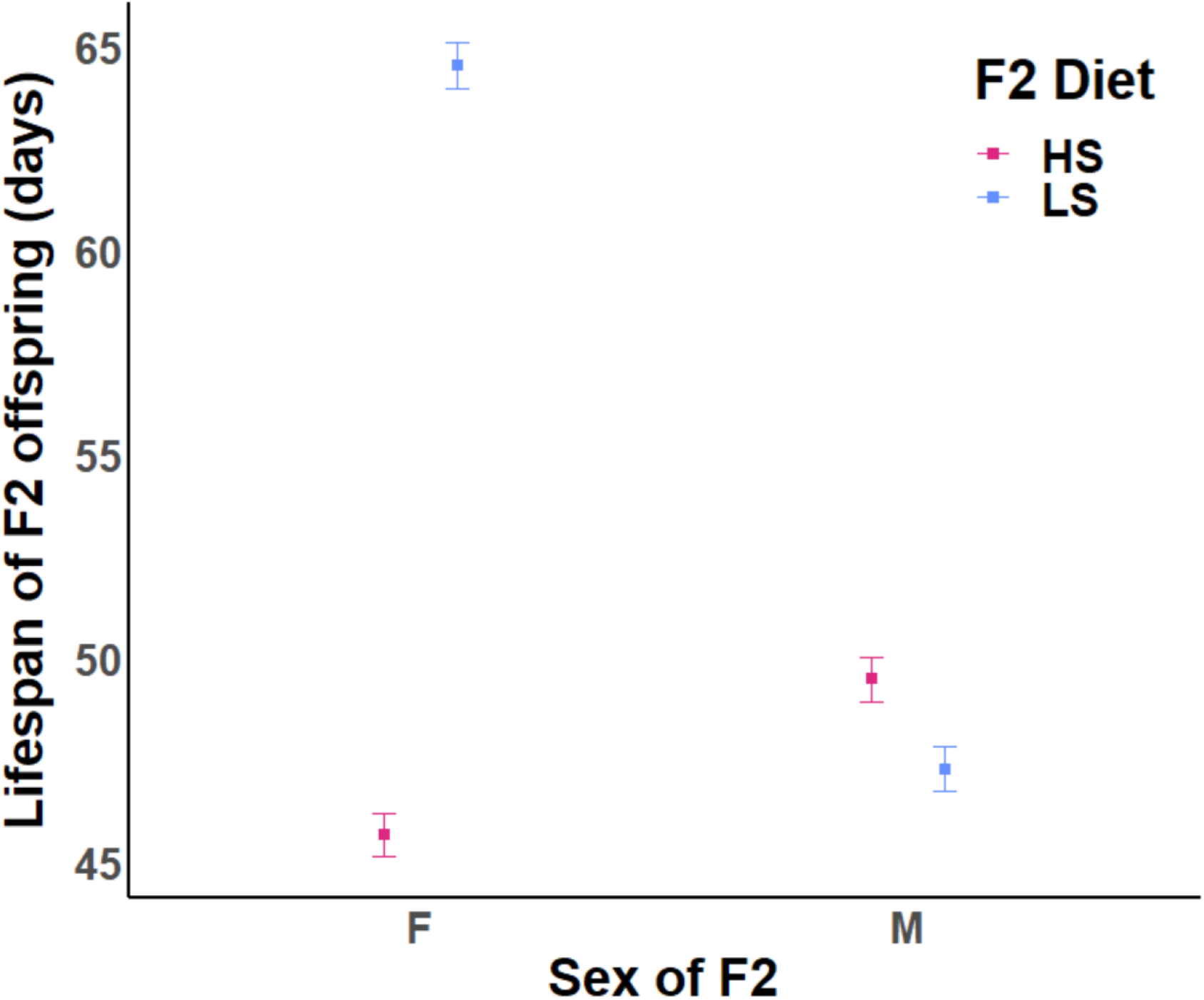
Direct effects of dietary sucrose on the lifespan (± standard error) of F2 granddaughters (F) and grandsons (M). HS indicates a high sucrose diet of 20% (P:C ratio 1:5.3), LS indicates a low sucrose diet of 2.5% (P:C ratio 1:1.4).

The lifespan of F2 flies was also in part mediated by the diets of their grandmothers, with the pattern of effects differing across F2 males and females (*F*_1,148_ = 9.35,*p* <0.01, Table S2, Figure 3, panel A). The transgenerational effects of sucrose concentration mimicked the direction of direct effects described above. That is, F2 females descended from grandmaternal lineages assigned to a low sucrose diet lived longer than those descended from high sucrose lineages, while the opposite pattern was observed in F2 males, whereby those descended from high sucrose grandmaternal lineages outlived those from low sucrose lineages (Figure 3, panel A). Additionally, matching combinations of grandmaternal-grandoffspring dietary sucrose led to shorter F2 lifespan than mismatched combinations (Figure 3, panel B).

**Figure 3.**
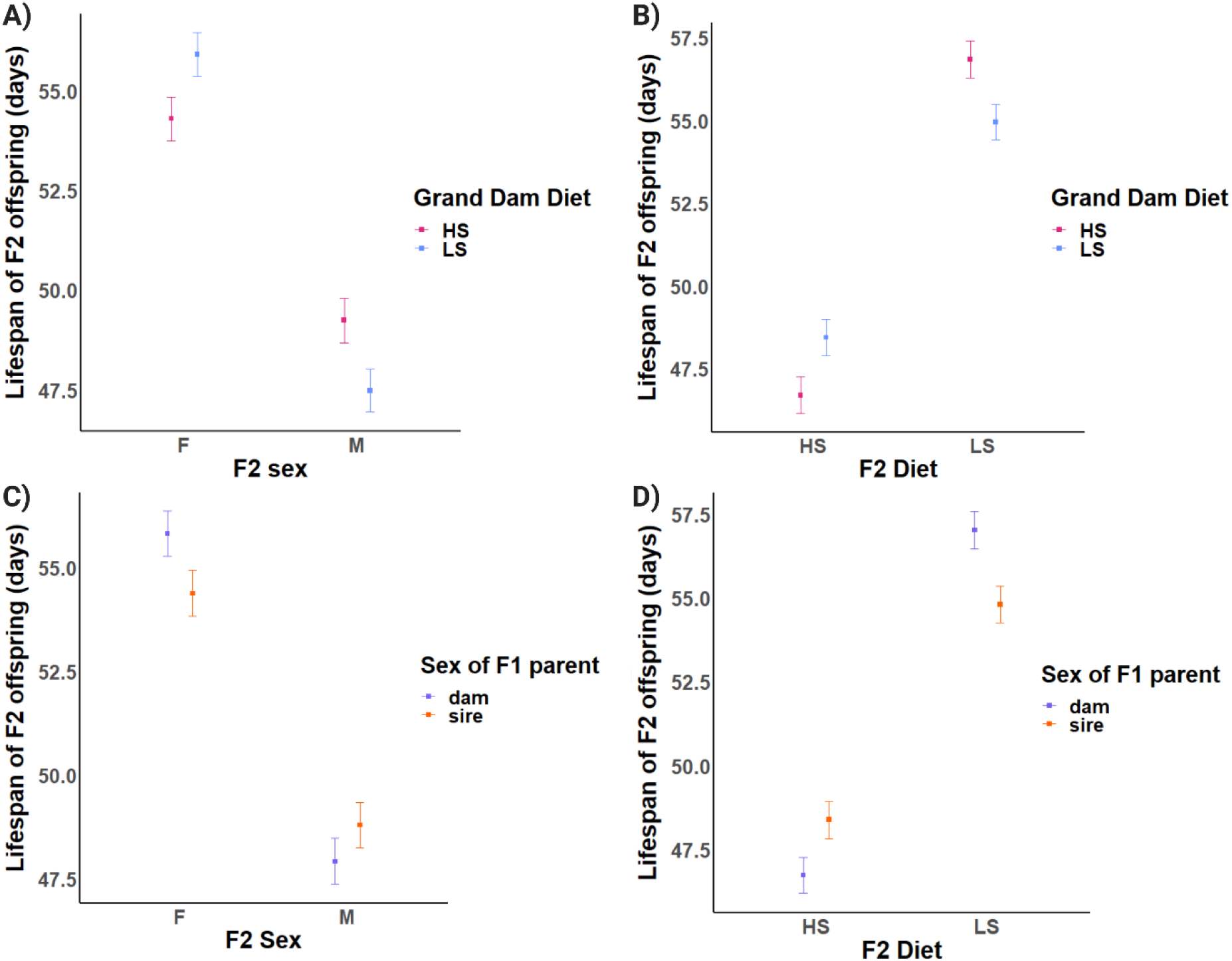
Effects of high sucrose (20% of overall solution) and low sucrose (2.5% of overall solution) on F2 lifespan. Plots show means, and standard error bars. (A) Lifespan of F2 flies (y-axis), their grand dam’s diet (colour), their sex (x-axis), (interaction: grand dam diet × F2 sex). (B) Lifespan of F2 flies (y-axis), their grand dam’s diet (colour), their diet (x-axis), (interaction: grand dam diet × F2 diet). (C) Lifespan of F2 flies (y-axis), the sex of the parental linage that received a diet treatment, (colour), their sex (x-axis), (interaction: F1 sex × F2 sex). (D) Lifespan of F2 flies (y-axis), the sex of the parental linage that received a diet treatment (colour), their diet (x-axis), (interaction: F1 sex × F2 diet).

In our experimental design, grandparental flies were manipulated, and F2 phenotypes measured. This involved transfer of effects across an intermediate generation – the F1 parents. Although the diets of F1 parents were never manipulated (they received a standard diet of 5% sucrose, an intermediate sucrose content), our experimental design ensured the grandparental effects were transferred through either male F1 or female F1 flies (but not both, Figure 1). Thus, we could track whether the sex of the *transferring F1 parents* affected the pattern and direction of the transgenerational effects. Indeed, the interaction between the sex of the F2 flies and the sex of the transferring F1 parents affected F2 lifespan (*F*_1,148_ = 4.44, *p* <0.05, Table S2); female F2 lived longer if the grandparental dietary treatments were transferred through F1 females, while male F2 lived longer when the effects were transferred through F1 males (Figure 3, panel C). The sex of the transferring F1 parent flies also moderated the direct effects of the F2 diet on F2 lifespan, Table S2, Figure 3, panel D, *F*_1,148_ = 12.42, *p* <0.001). F2 flies assigned directly to a high sugar diet lived longer if grandparental dietary treatments were transferred through F1males rather than through females, while F2 flies assigned to a low sugar diet lived longer if grandparental dietary treatments were transferred through F1 females than males.

### Grandoffspring fecundity, viability, and triglycerides are mediated by grand maternal and grand paternal diets

#### Fecundity & viability

Direct dietary effects were observed in the F2 generation; F2 Females had higher fecundity when ingesting the low sucrose than high sucrose diet. These direct effects of diet were, however, shaped by the grand paternal, but not grand maternal diet (Table S3, *F*_1_ = 5.49, *p* <0.05). Mismatched combinations of grandpaternal-F2 female diet resulted in F2 granddaughters producing more eggs than matched combinations (Figure 4, panel A). Female F2 fecundity was also shaped by an interaction between the grand maternal and grand paternal diets (Table S3, Figure 4, panel B, *F*_1_ = 14.77, *p* <0.05); F2 females that descended from matched grandmaternal-grandpaternal combinations tended to have lower fecundity than those arising from mismatched combinations, and in particular F2 females descended from grandparents that were each assigned to low sucrose diets exhibited lowest fecundity (Figure 4, panel B). The reproductive success (as gauged by the number of adult offspring produced) of the F2 females was also shaped by a similar interaction between grandmaternal and grandpaternal diet, in which the clutch size was lower for F2 females descended from matched, relative to mismatched, combinations of grandmaternal-grandpaternal diet (Table S4, Figure 4, panel C, *F*_1_ = 5.25,*p* <0.05).

**Figure 4.**
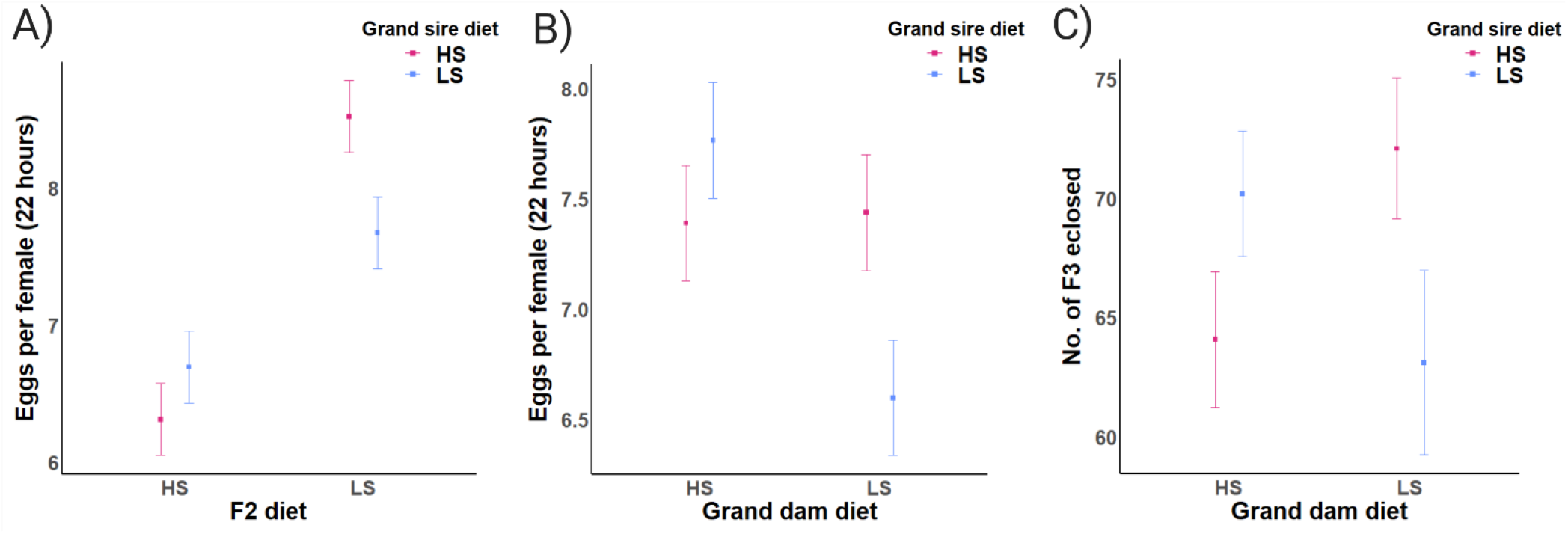
Effects of high sucrose (20% of overall solution) and low sucrose (2.5% of overall solution) on female F2 reproductive output. Plots show means, and standard error bars. (A) Number of eggs laid by F2 flies (y-axis), their grand sire’s diet (colour), their diet (x-axis), (interaction: grand sire diet × F2 diet). (B) Number of eggs laid by F2 flies (y-axis), their grand sire’s diet (colour), their grand dam’s diet (x-axis), (interaction: grand sire diet × grand dam diet). (C) Number of F3 flies eclosed per vial (y-axis), their grand sire’s diet (colour), their grand dam’s diet (x-axis), (interaction: grand sire diet × grand dam diet).

#### Triglyceride levels

An interaction between the diet of F2 offspring and the grandmaternal diet affected the triglyceride level of the F2 flies (Table S5, Figure 5, panel A, *F*_1,98_= 8.56,*p* <0.01). F2 flies fed high sucrose diets that descended from grandmothers assigned to high sucrose, exhibited much higher triglyceride levels than F2 flies from any other combination of grandmaternal-F2 offspring diet (Figure 5, panel A). Similarly, the interaction between F2 diet and grandpaternal diet shaped triglyceride level; however in this case, F2 offspring assigned to a high sucrose diet and descended from grandfathers assigned to low sucrose, exhibited much higher triglyceride levels than any other combination of grandpaternal-F2 diet (Table S5, Figure 5, panel B, *F*_1,98_ = 12.75, *p* <0.001).

**Figure 5.**
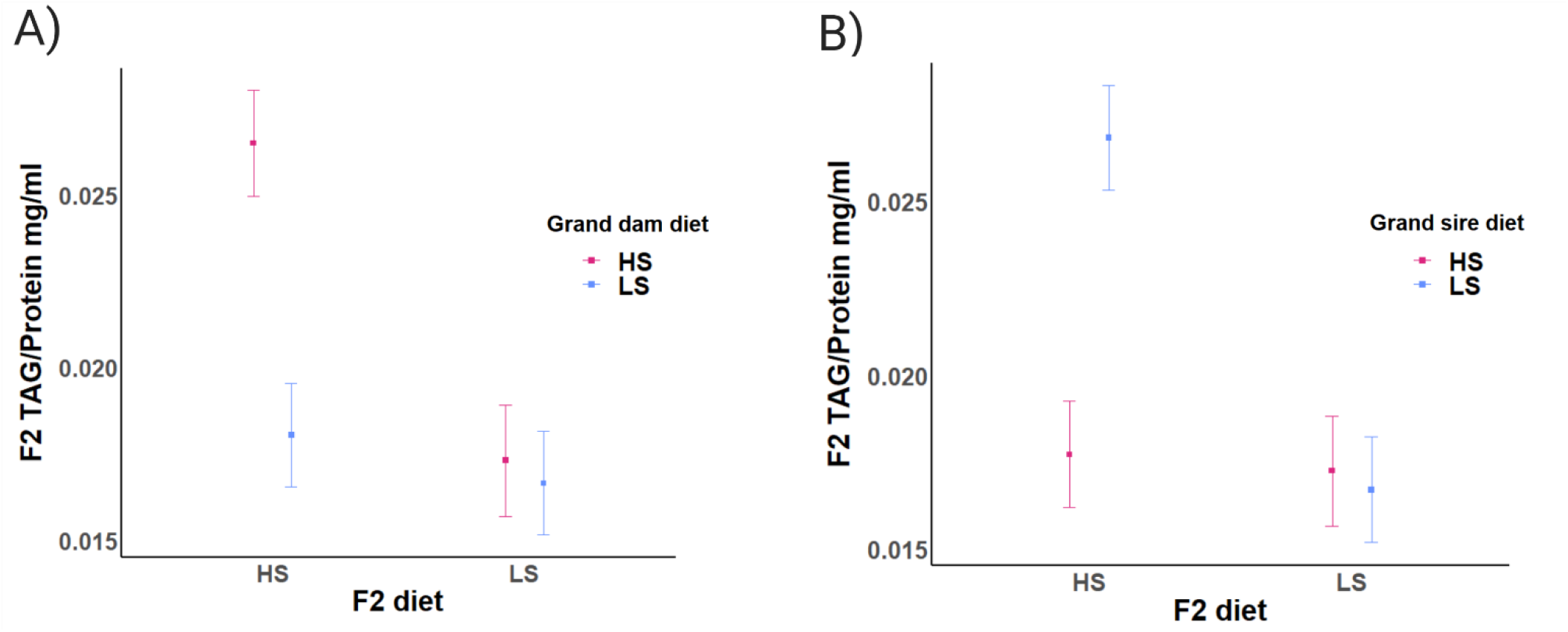
Effects of high sucrose (20% of overall solution) and low sucrose (2.5% of overall solution) on F2 whole body triglyceride (TAG) levels divided by their whole body protein levels, per fly. Plots show means, and standard error bars. (A) F2 TAG per fly (y-axis), their grand dam’s diet (colour), their diet (x-axis), (interaction: grand dam diet × F2 diet). (B) F2 TAG per fly (y-axis), their grand sire’s diet (colour), their diet (x-axis), (interaction: grand sire diet × F2 diet).

### No direct effect of dietary sucrose on female F0 fecundity

Neither the male, nor female diet, affected egg output of the F0 females (Table S1).

## Discussion

Here, we show opposing effects of dietary sucrose on the lifespan of each sex, in *D. melanogaster—*low sucrose enhances female lifespan, but decreases male lifespan relative to high sucrose. Notably, these effects were observed in both direct and indirect (i.e. transgenerational) contexts. Moreover, the dietary-mediated transgenerational effects on lifespan mimicked the observed direct effects for each sex: a low sucrose grandmaternal diet conferred elevated F2 female lifespan, but decreased male F2 lifespan, relative to a high sucrose grandmaternal diet. We also revealed strong effects of specific combinations of grandparental and grandoffspring diet, and between grandmaternal and grandpaternal diets, in shaping the measured traits; all of which exhibited a similar pattern—a mismatch in diet enhanced trait expression in subsequent generations. We highlight inherent complexity in the nature of the transgenerational effects. The effects are generally sex-specific, and unexpectedly, are affected by the sex of the transferring F1 parent. Finally, we note that interactions between grandparental diet and F2 diet affected triglyceride levels of F2 flies, suggesting that dietary-mediated modifications of triglyceride levels, across generations, may contribute to the observed transgenerational effects on life history phenotypes.

Studies investigating sex-specificity of transgenerational effects across a range of taxa have observed instances in which environmental modification such as dietary challenges, presence of predators, or behaviour-modifying drugs of the grandparental environment triggered sex-specific effects on grandoffspring phenotype. Intriguingly, in these cases, transgenerational effects tend to manifest in the opposite sex to that subjected to the grandparental treatment; that is, modification of the grandmaternal environment may enhance or inhibit trait expression among grandsons, or conversely, modification to the grandpaternal environment may enhance or inhibit trait expression among granddaughters ^10,25,30,32,33^. Our findings are consistent with previous research, revealing opposing directions of sucrose-mediated grandmaternal effects in each of the sexes. Notably, we have uncovered further levels of complexity in the nature of the transgenerational effects.

First, we revealed sex differences in the magnitude of transgenerational effect (the effect transmitted from dietary-treated F0 flies to F2 flies) are dependent on the sex of the transferring F1 parent. Second, we observed that the outcomes of transgenerational effects depend on interactions between the diets of the grandparents and those of the grandoffspring; dietary mismatching across generations tends to enhance lifespan (mediated by a grandmaternal-by-grandoffspring interaction) and fecundity (mediated by a grandpaternal by granddaughter diet interaction). Whether or not these effects are mediated by underlying triglyceride levels of the experimental flies remains unclear; yet one pattern was notable, suggestive of a possible transgenerational link between physiology and lifespan. F2 offspring assigned to a high sucrose treatment, and descended from high sucrose grandmaternal lineages exhibited the highest triglyceride levels and the shortest lifespans.

Investigations into life history traits are imperative in assessing the adaptive significance of transgenerational effects on offspring, given the close link between these traits and lifetime fitness^5^. Our experiments, across three generations, with the diet challenge also given to the F2 generation had the requisite power to address these previous knowledge gaps. Our finding that dietary mismatching (between both grandparents and between grandparents and grandoffspring) tends to enhance trait expression adds new insight to studies investigating transgenerational effects of diet, and of transgenerational effects of environmental change more generally. Previous studies of dietary-mediated transgenerational effects have tended to focus on changes in metabolite profiles and physiology across generations, rather than on changes to expression of life history traits^25,26^. Moreover, these designs typically do not have the requisite power to partition relative influences of (grand)maternal and (grand)paternal effects on transgenerational phenotypes, nor the factorial design required to determine whether transgenerational mismatches enhance or depress performance ^9^.

The prevailing prediction is that a matching of environment between grandparents and grandoffspring may augment offspring fitness-related traits, because the matching environments may allow parents to prime offspring to cope with environments that their parents faced (anticipatory effects). The evidence for anticipatory effects across contexts and taxa is, however, mixed and weak^9,29^, and many studies that have leveraged experimental designs with the power to test for these effects have primarily focused on intergenerational effects (from F0-F1^29^), with very few studies classified as transgenerational where grandoffspring should have no direct experience of the grandparental environment^34^. Our study generally revealed patterns that were contrary to the predicted pattern – dietary mismatching, rather than matching, between grandparents and F2 offspring tended to augment offspring performance. This begs the question of whether cross-generational dietary mismatching may be a general phenomenon that extends across the diets used in our study.

Two recent studies shed some light on this question. Deas et al. (2019) manipulated dietary quality across three generations (F0 to F2) in *D. melanogaster*, providing flies of each generation with a ‘rich’ diet (rich in calories and supplemented with yeast) or a poor diet (calorie diluted, with no yeast supplementation), in all combinations, and then measuring phenotypic expression in the grandoffspring (F2). They reported that a mismatch between the diet quality (“poor vs “good” diet) of granddams and granddaughters led to a faster development time in the pupal stage of the granddaughters, but this effect did not hold for the entire development time^35^. This study also focused on females, and therefore was not able to capture sex specificity in any generation. On the other hand, Camilleri et al. (2022) tested effects of dietary mismatching of F0 flies and their F1 offspring, manipulating the diets of parents of each sex and their offspring, and utilising the same sucrose diets used in the current study. We found that dietary mismatching between parents and F1 offspring led to an increase in lifespan, and fecundity of the offspring^28^. Here, we advance these findings by demonstrating that these effects of dietary mismatch are carried over for multiple generations, are also dependant on the sex of F1 lineage. Because the effects are unambiguously transgenerational (extending from F0 to F2), they are less likely to result from differences in condition of the grandparents, suggesting instead possible epigenetic mechanisms regulating the effects.

In sum, our work uncovers dietary-mediated transgenerational effects that are on the one hand remarkably consistent across generations – transgenerational effects of sucrose tend to mimic the direct effects. We have also extended previous work to demonstrate that dietary mismatching across generations tends to augment phenotype in a manner that is unlikely to be directly linked to condition-dependence. Future work should focus on uncovering the ecological and evolutionary significance of these results, and the underpinning mechanistic drivers. We suggest that a process in which transgenerational dietary mismatching promotes fitness of future generations could buffer populations from future changes in environment, and be particularly adaptive for species that live and depend on ephemeral resources for their source of nutrients. If this is the case, then populations evolving in fluctuating environments may be more likely to evolve mechanisms that promote the fitness of offspring encountering novel environments.

## Methods

### Study species and generating experimental flies

We sourced flies from Dahomey, a large laboratory population of *D. melanogaster*, originally sourced from Benin West Africa^36^. The flies have been maintained in large population cages, with overlapping generations in the Piper laboratory, Monash University, Australia, since 2017, and prior to that in the Partridge laboratory, University College London ^37^. Prior to the beginning of the experiment, we collected ~3000 eggs from the cages, and distributed them into 250mL bottles containing 70mL of food. Food comprised 5% sucrose (50 grams sucrose, 100 grams yeast, 10 grams agar per 1 litre solution with an estimated protein to carbohydrate [P:C] ratio of 1:1.9, and 480.9 kcal per litre (see Supplementary Material Figure S4 for further diet details). Every generation (for 7 generations), adult flies eclosing from multiple bottles were admixed prior to redistributing the flies across new bottles. To control for potential sources of variation in their environment, during these 7 generations we strictly controlled both the age of flies at the time of ovipositioning—all flies were within 24 h of eclosion into adulthood when producing the eggs that propagated the subsequent generation, and their population density was 300-320 adult flies within each bottle in each generation.

### Dietary treatments

The diet media we used consists of sucrose, autolysed brewer’s yeast powder (sourced from MP Biomedicals SKU 02903312-CF), and agar (grade J3 from Gelita Australia), as well as preservatives—propionic acid, and nipagin. We prepared two dietary treatments, differing in relative sucrose concentration; 2.5% sucrose (that we refer to as a lower sucrose treatment relative to the 5% concentration usually provided to the population of flies used in this experiment), and 20% sucrose (that we refer to as a higher sucrose treatment) of overall food solution. The 2.5% sucrose diet contains 25 grams of sucrose, 100 grams of yeast and 10 grams of agar per litre of food prepared, with an estimated P:C ratio of 1:1.4 and 380.9kcal per litre of food. The 20% sucrose treatment contains 200 grams of sucrose, 100 grams of yeast, and 10 grams of agar per litre of food prepared, with an estimated P:C ratio of 1:5.3 and 1080.9kcal per litre of food. The diets thus differed not only in sucrose concentration, but overall macronutrient balance and their total caloric content, resembling differences typically observed between obesogenic and healthy diets in humans. The higher sucrose concentration was selected based on preliminary experiments that we conducted, and which elicited an obese-like phenotype in the flies, consistent with results from previous work in *D. melanogaster* ^11,25,38^. All diets contained 3ml/l of propionic acid and 30ml/l of a Nipagin solution (100g/l methyl 4-hydroxybenzoate in 95% ethanol) and were cooked according to the protocol described in Bass *et al.* (2007) ^39^. Each vial is 40mL in volume, and contained 7mL of food.

### Experimental design

Male and female virgin flies were assigned to one of two of the dietary treatments prior to mating (we refer to this generation of flies as F0), and then the grandoffspring produced (F2 generation) were also assigned to one of the two treatments. All possible combinations of grand dam × grand sire × grandoffspring diet treatment were represented (= 2 × 2 × 2 = 8 combinations). Specifically, we collected 1280 flies of the F0 generation as virgins and placed them onto either the high sucrose (20%) or the low sucrose (2.5%) diets for the first 6 days of their adult life. They were in vials of 10 flies across 64 vial replicates per treatment, and per sex (High sucrose: 32 vials of males and 32 vials of females; low sucrose, 32 vials of males and 32 vials of females, 128 vials in total;1280 flies, 640 of each sex). They were kept in their respective sexes. We transferred flies to vials containing fresh food of the designated diet every 48 hours during this 6 day period.

At day 6, we randomly sampled six vials from each treatment, and snap froze (using liquid nitrogen) the flies of these vials, storing them at −80°C for subsequent measures of triglyceride levels. Cohorts of flies in the remaining vials then entered a cohabitation phase to enable female and male F0 flies to mate. Cohorts of males and female flies were combined, in vials of 10 pairs, in each of all four possible diet combinations: lower sucrose females × lower sucrose males; higher sucrose females × higher sucrose males; lower sucrose females × higher sucrose males; higher sucrose females × lower sucrose males. During this phase, flies cohabited for 96 hours. They were transferred to a new vial with fresh food of standard 5% sucrose diet every 24 hours during this time.

The vials from the 6 day old F0 flies (i.e., the vials from Day 1 of the 96 h cohabitation phase) were retained, and the eggs that had been laid by females of the respective vials were trimmed to 80 per vial by removing excess eggs with a spatula. The remaining eggs were left to develop into adult offspring over 10 days at 25°C (on a 12:12 light/dark cycle in a temperature-controlled cabinet; Panasonic MLR-352H-PE incubator). These adult flies constituted the F1 offspring in the experiment, and F1 flies developed on standard 5% sucrose media. We collected 2080 virgin F1 flies from each of the four combinations of parental diet treatments, and placed them in sex-specific cohorts of 10 individuals per vial, on standard 5% sucrose media for 6 days. We then allowed these F1 males and F1 females to cohabit and mate with male or female *tester* flies (creating 10 pairs per vial) that had been collected from the same Dahomey stock population (but not subjected to a dietary sucrose treatment) to create the F2 generation. The diet treatments applied to the F0 flies were thus transferred to the F2 generation via either F1 males or F2 females, but never through both sexes. The F1 flies were 6 days of adult age when laying the eggs that produced the F2 generation.

We then collected virgin F2 flies – the grandoffspring of the F0 flies – from each of the four combinations of F0 diet treatments (per sex), and placed them in their respective sexes in vials of 10 flies, across vial replicates per treatment per sex (4080 flies, 2040 male, 2040 female). We then assigned these F2 flies, produced by each dietary treatment combination of F0 flies, to either the lower sucrose or higher sucrose diet. At day 6 of adulthood, we snap froze F2 flies of six randomly chosen vials per grand dam × grand sire × grandoffspring combination. On the same day, 10 virgin focal F2 flies of each grand dam × grand sire × grandoffspring combination and each sex were placed together with 10 age-matched tester flies of the opposite sex from the Dahomey population, entering into a cohabitation phase of 96 h (during which time the number of eggs laid by females of each vial was assessed). After 96 hours flies were separated again into their respective sexes (in vials of 20 flies), and assigned back onto either the lower sucrose or higher sucrose diets that they had been on prior to cohabitation, and a lifespan assay carried out.

### Lifespan

We scored the lifespan of experimental flies of the F2 generation. Each vial in the assay commenced with 20 flies of single sex in each, and we included 10 vial replicates per treatment (grand dam × grand sire × grandoffspring) (3400 flies total, the original amount collected, minus the snap frozen samples). The number of dead flies per vial was scored three times per week (Monday, Wednesday, Friday), and surviving flies at each check transferred to vials with fresh food of the assigned diet treatment—until all flies were deceased. During the lifespan assay, vials were stored in boxes (of 85 vials per box) that were moved to randomised locations in a (25°C) control temperature cabinet every few days to decrease the potential for confounding effects of extraneous sources of environmental variation within the cabinet from affecting the results.

### Fecundity

We measured the egg output of female flies from generations F0 and F2 at eight days following eclosion, as a proxy of female fecundity. On day eight, female flies oviposited for a 23 hour period, and were then transferred to fresh vials. Day eight was selected because fecundity over 24 hours at this age has been shown to correlate with total lifetime fecundity of females in this Dahomey population^39^ and early, short term measures of reproduction of between one and seven days can be used to accurately predict total lifelong fecundity in *D. melanogaster*^40^. Moreover, data shows that varying the range of sucrose concentrations did not alter the timing reproductive peaks between treatments ^39^.

For the F0 generation, we counted eggs from vials, each containing 10 female flies, that had been mated with 10 male flies, across 2 different sucrose levels (2.5% and 20% sucrose), and different mate combinations, as above. For the F2 generation, we counted eggs from each grand dam × grand sire × grandoffspring dietary treatment combination; each combination was represented by 10 vial replicates, each containing 10 focal females (females from the experiment) combined with 10 tester male flies. Additionally, we counted the number of adult flies that eclosed within 10.5 days from the eggs laid by F2 females (a composite of clutch viability and juvenile developmental speed). F2 females cohabited and mated with age-matched tester males of the Dahomey population (in the experimental process described above, rather the standard medium of 5% sucrose), for 24 hours at 6 days of life, and the vials containing these eggs were left to develop into adult offspring, for 10 days at 25°C; 12:12 light/dark cycle in a temperature-controlled cabinet (Panasonic MLR-352H-PE incubator).

### Lipids and protein

Whole-body triglyceride levels were measured in adult flies from the F2 generation (six days of adult age, corresponding with six days of exposure to the relevant F2 dietary treatment, prior to mating) and normalized to protein content (full protocols reported in the Supplementary Material). Three biological replicates per treatment level, with three technical replicates per biological replicate were used. Five female flies and eight male flies respectively, were used for each biological replicate in the assay.

### Statistical Analyses

We used R (Version 3.6.1) and RStudio (Version 1.2.1335) (R Core Team, 2019) for statistical analyses. To test the effects of F0 female diet, F0 male diet, F2 diet, and sex on lifespan, TAG, and F2 offspring production, we fitted linear mixed effects models, using the R package lme4^41^, to the lifespan data for the F2 generation. We use the term lifespan to denote the age of recorded death for each individual fly within a margin of 72 hours (for example, a lifespan of 30 days indicates that a fly died between 27-30 days post eclosion). To test the effects of grand maternal diet, grand paternal diet, grand-offspring diet, and sex on female fecundity, we fit a linear model to the egg output data for both generations.

We included F0 male, F0 female, F2 diets, and F2 sex as fixed effects in each model, exploring interactions between these factors. We included the vial identification number as a random effect in the lifespan models. The fecundity models only included one observation per vial because we counted eggs per vial, and divided by the number of females in the vial (approx. 10 females); therefore no random effects were included in this model. We used log-likelihood ratio tests that reduce the full model, via the sequential removal of highest order terms that did not (significantly) change the deviance of the model, using a *p* value significance level of <0.05. The final reduced models (except fecundity measures) were fit by restricted maximum likelihood, applying type III ANOVA with Kenwood-Roger’s F test and approximation of denominator degrees of freedom. We used sum to zero constraints in all models, and we visually inspected diagnostic plots for the linear mixed effect models, to ensure that the assumptions of normality and equal variances were met.

## Supporting information

All supplemental information

## Author contributions

TLC, DKD, MDWP & RLR designed the experiment, TLC planned and carried out the experiment, and wrote the initial draft of the manuscript, and TLC, DKD, MDWP & RLR all contributed to the writing and editing of the manuscript. TLC performed statistical analysis under the guidance of DKD.

## Acknowledgements

The authors are grateful for the help they received in the laboratory from: Pavani Manchanayake, James Wang, Skye Bulka, Natalie Wagan, Rebecca Koch, and Winston Yee. Indispensable guidance with molecular work was provided from Amy Dedman. Additional invaluable assistance was provided by Caleb Carter.

