## Supplementary material for "Sex-specific transgenerational effects of diet on offspring life history and physiology": All supplemental information

### Supplementary information: Results

#### Tables

**Table S1.** Statistical results (analysis of deviance) of the linear model (Gaussian error distribution) after model reduction for predictors of female F0 egg output.
*** indicates significance at $p < 0.001$, ** indicates significance at $p < 0.01$, * indicates significance at $p < 0.05$.

| ***Fixed Effects*** | ***Sum Sq*** | ***F*** | ***p-value*** |
|---|---|---|---|
| (Intercept) | 1143.66 | 364.07 | < 0.001 *** |
| Male diet | 8.97 | 2.979 | 0.09117 |
| Female diet | 0.64 | 0.2155 | 0.6539 |
| Residual | 135.53 | 45 | |

All *df* =1

**Table S2.** Statistical results (analysis of deviance with Kenward-Roger method) of the linear mixed model (Gaussian error distribution) after model reduction for predictors of F2 offspring age.
*** indicates significance at $p < 0.001$, ** indicates significance at $p < 0.01$, * indicates significance at $p < 0.05$.

| ***Fixed Effects*** | ***F*** | ***Df.res*** | ***p-value*** |
|---|---|---|---|
| Intercept | 2639.15 | 136.92 | < 0.001 *** |
| F2 diet | 430.37 | 145.66 | < 0.001 *** |
| F2 sex | 15.73 | 147.58 | < 0.001 *** |
| Grand maternal diet | 13.16 | 145.25 | < 0.001 *** |
| Parental (F1) sex | 0.2757 | 146.11 | 0.6003 |
| F2 diet : F2 sex | 369.80 | 148.33 | < 0.001 *** |
| Grand maternal diet : F2 sex | 9.35 | 148.35 | < 0.01 ** |
| F2 diet : grand maternal diet | 11.01 | 148.44 | < 0.01 ** |
| F2 sex : F1 sex | 4.44 | 148.47 | < 0.05 * |
| F2 diet : F1 sex | 12.42 | 148.56 | < 0.001 *** |
| ***Random Effects*** | ***Variance*** | | |
| Vial identification | 1.423 | | |
| Residual | 12.73 | | |

*df* =1

**Table S3.** Statistical results (analysis of deviance) of the linear model (Gaussian error distribution) after model reduction for predictors of female F2 offspring egg output.
*** indicates significance at $p < 0.001$, ** indicates significance at $p < 0.01$, * indicates significance at $p < 0.05$.

| ***Fixed Effects*** | ***Sum Sq*** | ***F*** | ***p-value*** |
|---|---|---|---|
| (Intercept) | 1053.44 | 383.39 | < 0.001 *** |
| F2 fem diet | 97.71 | 35.56 | < 0.001 *** |
| Grand maternal diet | 0.05 | 0.02 | 0.8973 |
| Grand paternal diet | 13.04 | 4.75 | < 0.05 * |

| | | | |
|---|---|---|---|
| F2 fem diet : grand paternal diet | 15.08 | 5.49 | < 0.05 * |
| Grand maternal diet : grand paternal diet | 14.77 | 5.38 | < 0.05 * |
| Residual | 423.15 | | |

*df* =1, *Res df*=154

**Table S4.** Statistical results (analysis of deviance with Kenward-Roger method) of the linear mixed model (Gaussian error distribution) after model reduction for predictors of female grand offspring (F3) offspring viability (offspring produced by F2 females).
*** indicates significance at $p < 0.001$, ** indicates significance at $p < 0.01$, * indicates significance at $p < 0.05$.

| ***Fixed Effects*** | ***F*** | ***Df.res*** | ***p-value*** |
|---|---|---|---|
| Intercept | 321.888 | 5.754 | 0.20190 |
| Grand maternal diet | 4.145 | 33.062 | < 0.05 * |
| Grand paternal diet | 2.826 | 151.714 | 0.09479 |
| F2 fem diet | 31.59 | 107.955 | < 0.001 *** |
| Grand maternal : Grand paternal diet | 5.255 | 9.464 | < 0.05 * |
| ***Random Effects*** | ***Variance*** | | |
| Counter | 0.00 | | |
| Residual | 252.8 | | |

*df* =1

**Table S5.** Statistical results (analysis of deviance with Kenward-Roger method) of the linear mixed model (Gaussian error distribution) after model reduction for predictors of F2 whole-body TAG divided by protein.
*** indicates significance at $p < 0.001$, ** indicates significance at $p < 0.01$, * indicates significance at $p < 0.05$.

| ***Fixed Effects*** | ***F*** | ***Df.res*** | ***p-value*** |
|---|---|---|---|
| Intercept | 180.64 | 99.614 | < 0.001 *** |
| F2 diet | 3.32 | 98.185 | 0.064 |
| F2 sex | 2.25 | 96.717 | 0.136 |
| Grand maternal diet | 21.84 | 99.352 | < 0.001 *** |
| Grand paternal diet | 24.74 | 99.352 | < 0.001 *** |
| F2 diet : Grand mat diet | 8.56 | 98.832 | < 0.01 ** |
| F2 diet : Grand pat diet | 12.75 | 98.848 | < 0.001 *** |
| ***Random Effects*** | ***Variance*** | | |
| Plate reading replicate | 0.0031018 | | |
| Technical replicate | 0.0015873 | | |
| Vial identification | 0.0063383 | | |
| Residual | 0.0007295 | | |

All *df* =1

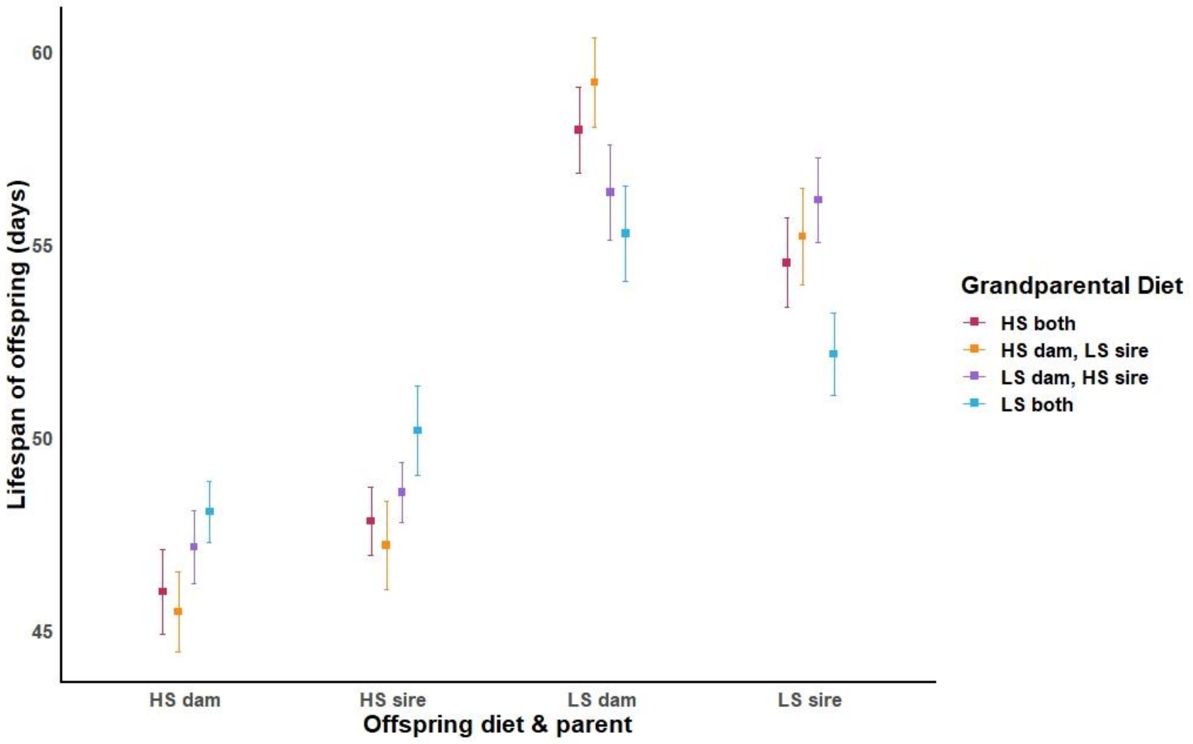


**Figure S1.** Mean longevity ± Standard Error (95% Confidence Interval) for the lifespan of the F2 generation. The dietary effects in the F0 generation (grandparental diets) were transferred to the F2 offspring either via F1 males or F1 females (but never via both sexes).

This figure the combination of the F2 and F1 combinations as well as the F2 combinations. HS indicates a high sucrose diet of 20% (P:C ratio 1:5.3), LS indicates a low sucrose diet of 2.5% (P:C ratio 1:1.4).

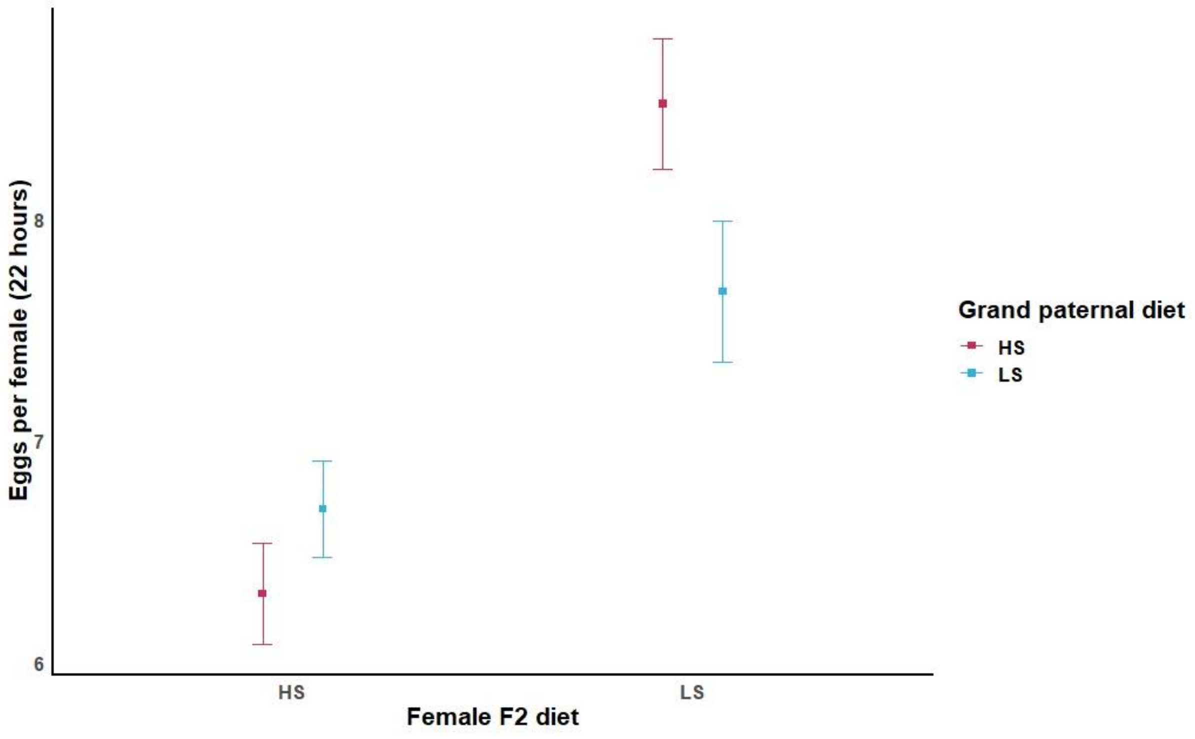


**Figure S2.** Mean egg output per female ± Standard Error (95% Confidence Interval) for the F2 generation, showing the grand paternal diet and the diet of the female F2 offspring. HS indicates a high sucrose diet of 20% (P:C ratio 1:5.3), LS indicates a low sucrose diet of 2.5% (P:C ratio 1.1.4).

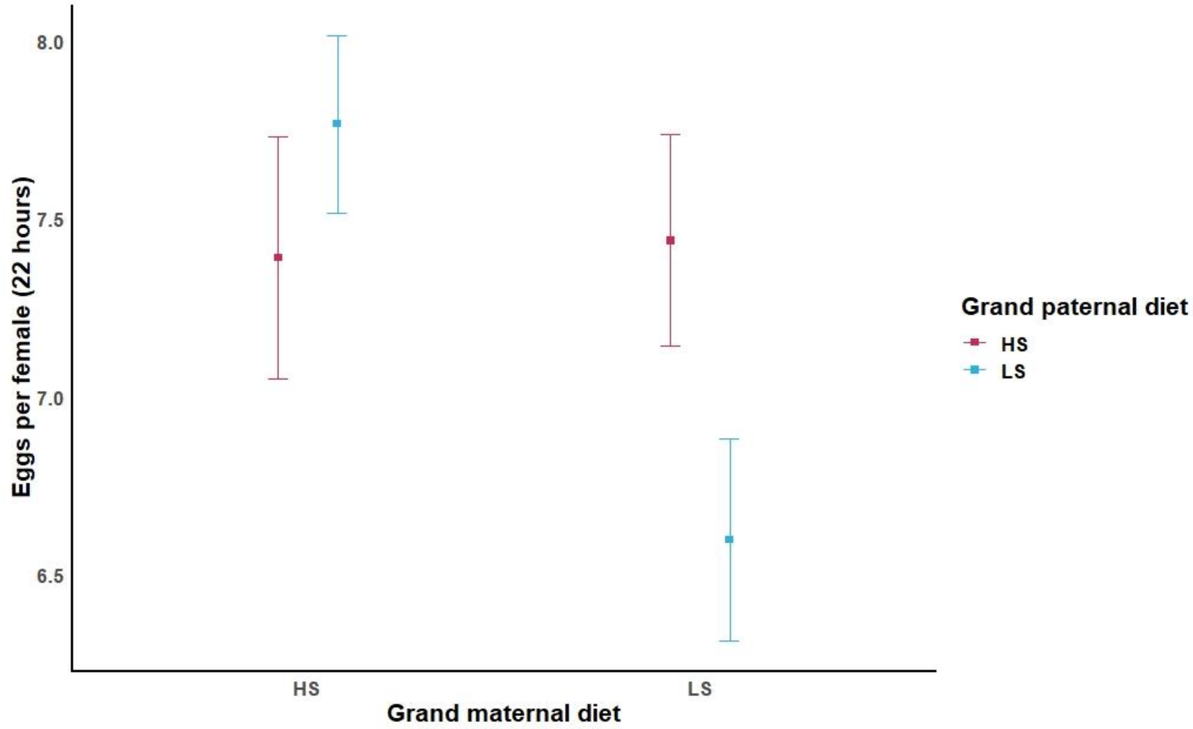


**Figure S3.** Mean egg output per female ± Standard Error (95% Confidence Interval) for the F2 generation, showing the grand paternal diet grand maternal diet. HS indicates a high sucrose diet of 20% (P:C ratio 1:5.3), LS indicates a low sucrose diet of 2.5% (P:C ratio 1:1.4).

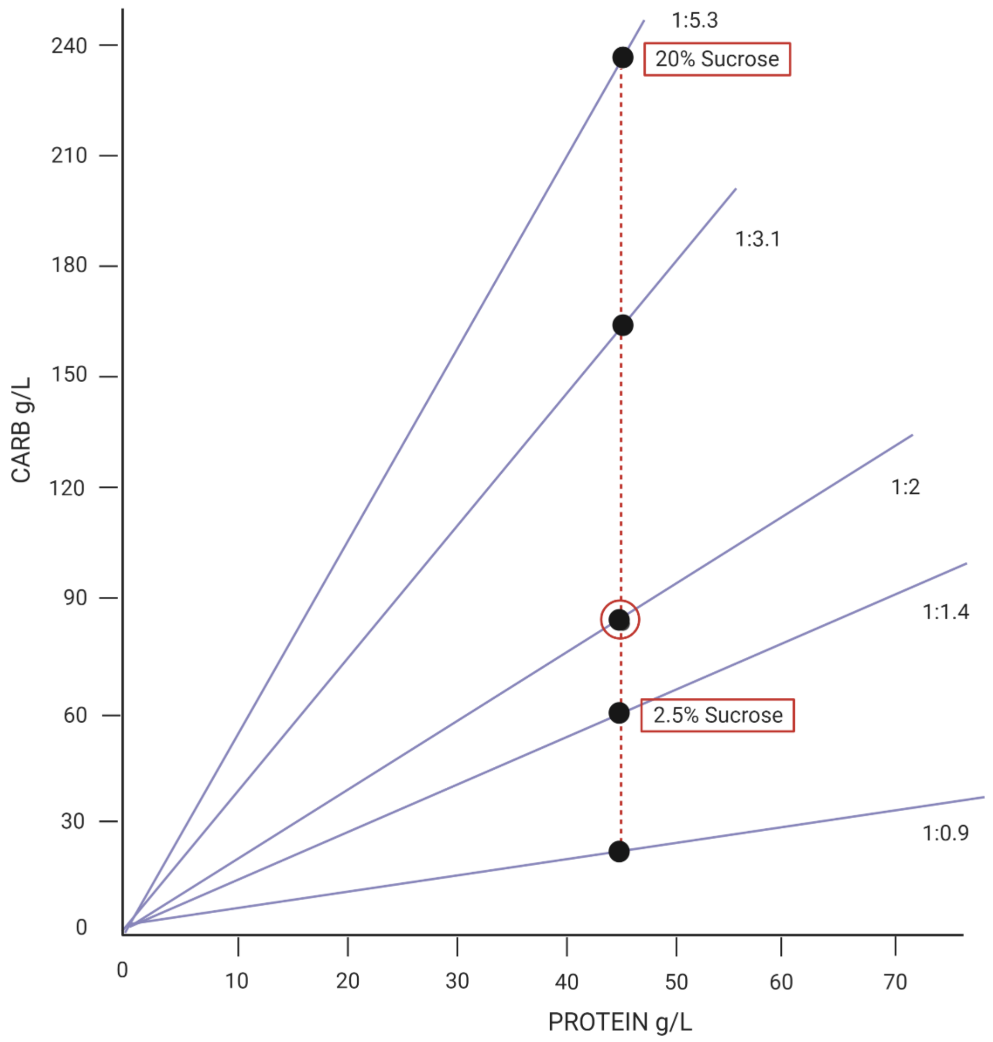


**Figure S4.** The diets used in the experiment according to their carbohydrate and protein contents, the diet in the middle that is circled represents the standard media in which flies were reared on prior to the experiment, while they were mating and for the full durations in the case of the F1 generation. The 20% sucrose is what we refer to as higher relative sucrose, and 2.5% sucrose we refer to as lower relative sucrose treatments.
